# A transcriptionally distinct subpopulation of healthy acinar cells exhibit features of pancreatic progenitors and PDAC

**DOI:** 10.1101/2021.01.04.425042

**Authors:** Vishaka Gopalan, Arashdeep Singh, Farid Rashidi, Li Wang, Eytan Ruppin, Efsun Arda, Sridhar Hannenhalli

## Abstract

Pancreatic ductal adenocarcinoma (PDAC) tumors can originate either from acinar or ductal cells in the adult pancreas. We re-analyze multiple pancreas and PDAC single-cell RNA-seq datasets and find a subset of non-malignant acinar cells, which we refer to as acinar edge (AE) cells, whose transcriptomes highly diverge from a typical acinar cell in each dataset. Genes up-regulated among AE cells are enriched for transcriptomic signatures of pancreatic progenitors, acinar dedifferentiation, and several oncogenic programs. AE-upregulated genes are up-regulated in human PDAC tumors, and consistently, their promoters are hypo-methylated. High expression of these genes is associated with poor patient survival. The fraction of AE-like cells increases with age in healthy pancreatic tissue, which is not explained by clonal mutations, thus pointing to a non-genetic source of variation. We also find edge-like states in lung and liver tissues, suggesting that sub-populations of healthy cells across tissues can exist in pre-malignant states.

## Introduction

Pancreatic ductal adenocarcinoma (PDAC) is one of the deadliest cancers with ∼8% survival rate at 5 years^1^. Pathogenesis of PDAC, and in particular, the cell of origin for PDAC, is not yet fully resolved, thus impeding development of robust therapies. Recent work has demonstrated that in mice, PDAC tumors can be driven from both acinar and ductal cells^2^, where an acinar to PDAC transformation is mediated by acinar-ductal metaplasia (ADM)^3^.

A classical view of cancer posits that oncogenesis is mediated by a series of somatic mutations in key oncogenes and tumor suppressors, accompanied by clonal selection^4,5^. While this clonal genetic model is widely accepted as one of the dominant pathways to oncogenesis, epigenetic alterations also play a key role. Indeed, transcriptional and epigenetic heterogeneity in the progenitor cell population forms the basis for later malignant transformation^6^, where such heterogeneity has been shown to be crucial for pre-malignant pancreatic lesions to progress to PDAC^7,8^. Furthermore, in a clonal cellular population, pervasive transcriptional fluctuation, in conjunction with complex regulatory networks, can result in a distinct meta-stable cellular states^9–11^. For instance, in a clonal population of blood progenitors, high SCA1-expressing outlier cells preferentially commit to the myeloid lineage, whereas cells with low SCA1 expression commit to proerythrocytes^10^. Taken together, this suggests a potential non-genetic basis for early stages of tumorigenesis, driven by transcription fluctuation across clonal cells resulting in a distinct cell state primed for malignant transformation in the favorable environment^11^. Oncogenic mutations can further amplify this non-genetic heterogeneity, as seen in breast epithelial cell cultures where oncogenic mutations increase the rate of switching between non-stem and stem-like epithelial cells^12^. An interplay between genetic and epigenetic alterations is likely to underlie complete malignant transformation^13^.

In this work, we investigated the potential role of transcriptional heterogeneity in pancreatic epithelial cells in priming PDAC. We analyzed a published single-cell transcriptomic dataset comprising 57,730 cells from 24 PDAC tumors and 11 pancreas samples from patients having non-PDAC indications^14^. We found that non-malignant acinar cells contained a sub-population, which we refer to as edge cells (following the terminology in Li et. al^15^), whose transcriptomes diverge from the average acinar cell and show features of pre-malignancy. In particular, genes that are differentially up-regulated among the acinar edge cells are enriched for transcriptomic signatures of pancreatic progenitors and acinar dedifferentiation, as well as several oncogenic programs such as Kras signaling, fatty acid metabolism, and epithelial-mesenchymal transition (EMT). Furthermore, in human PDAC tumors, the genes up-regulated in acinar edge cells are up-regulated and consistently, their promoters are hypo-methylated. Higher expression of these genes also associates with PDAC patient survival. This suggests potential clinical relevance of these early malignancy priming events in acinar cells. Finally, we validate the existence of acinar edge cells in additional independent pancreatic datasets and additionally find that the fraction of edge-like cells increases with age in healthy pancreatic tissue, thus providing a potential mechanism linking the known increase of PDAC incidence with age^1^. Intriguingly, we see strong functional similarity between transcriptional drift from non-edge to edge acinar cells and those previously reported in healthy to pre-malignant lung transformation^16^, suggesting that our observations in PDAC may possibly be more general. Indeed, beyond the pancreas, we found edge-like states among epithelial cells in non-malignant lung and liver tissues.

Overall, our work suggests that transcriptional heterogeneity among non-malignant epithelial cells may be large enough for a fraction to exist in a dedifferentiated, pre-malignant state. Since genes up-regulated in this pre-malignant state also increased in expression with age, this may help explain the higher incidence rate of tumors with age in these tissues, in addition to other putative mechanisms associated with the increase in cancer risk with aging^17^.

## Results

### Normal acinar cells include a transcriptionally divergent *Edge* subpopulation shifted toward a malignant state

We obtained processed gene-wise read counts from RNA-seq profiling of 57,730 pre-annotated cells across 24 PDAC and 11 non-PDAC samples^14^. The non-PDAC samples were taken from the normal pancreatic sites (Table S1 in Peng et. al^14^) of patients with other conditions: neuroendocrine tumors (n=3), solid pseudopapillary tumors (n=3), serous cystic neoplasia (n=1), mucinous cystic neoplasia (n=2), duodenal intraepithelial neoplasia (n=1) and small intestine papillary adenocarcinoma (n=1). We processed the data using Seurat v3.0^18^, following which we used doubletFinder^19^ to discard 2,877 potentially doublet cells, leaving us with 54,853 cells. These cells comprised 10 annotated types --T cells, B cells, Macrophages, Stellate cells, Fibroblasts, Endothelial cells, Acinar cells, Ductal cells (Type 1 and Type 2) and Endocrine cells. A UMAP plot of the data shows that the annotated cell types are well-separated (Fig. S1). In the original annotations of the data, Ductal cell type 2 refers to malignant ductal cells, to contrast them with non-malignant ductal cells (type 1).

If a given non-malignant cell cluster, say X, passes the two statistical filters below, we state that X contains edge cells (Figures 1A). The first filter – heterogeneity test – checks if a subset of cell in X have significantly diverged from X’s medoid in Principal Component (PC) space. These PCs, which we call Normal PCs, are computed based on transcriptomes only in X to capture gene expression variation within X. If X passes the filter, we consider the 10% of cells farthest from X’s medoid as candidate edge cells. The second filter – proximity test – checks if the candidate edge cells are significantly closer to the malignant cluster than the remaining cells in X. The proximity test is based on PC coordinates computed from cells in both X and the malignant cluster, which we call Pooled PCs. Technically, the heterogeneity test can also be carried out in Pooled PC space. However, since Pooled PCs also capture gene expression differences between X and the malignant cluster, they do not provide an unbiased measure of heterogeneity within X.

**Fig. 1.**
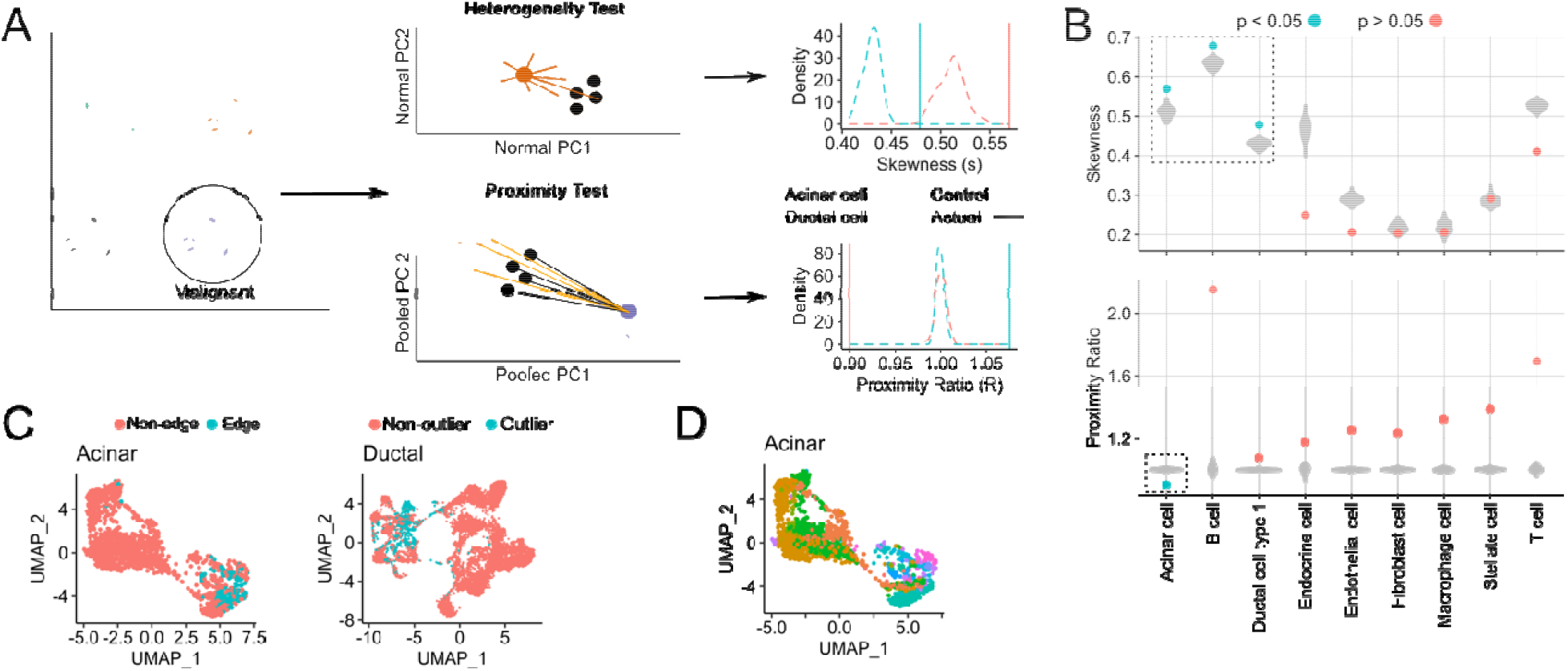
Testing the presence of an edge sub-population among non-malignant cells in scRNA-seq data. (A) Within each non-malignant cluster, every cell’s distance from the cluster medoid (in Normal PC space) is calculated, and the resulting distance distribution is tested for positive skewness. In the proximity test, we test if the 10% of non-malignant cells farthest from their own medoid (black, termed outlier cells) are significantly closer, in the Pooled PC space, to the malignant cluster medoid (dark purple) than the remaining 90% of cells (orange). If both test conditions hold, the outlier cells are called edge cells. For both tests, examples of the distributions of skewness and the proximity ratio are shown for acinar and ductal cells, as well as their respective control populations (B) Violin plots of medoid distance distribution skewness values (top) and malignant proximity ratio (bottom) after shuffling is performed 100 times for each indicated cluster. Filled circles indicate skewness and proximity ratio values of actual cells, where blue and red indicate a significant (< 0.05) or insignificant p-value for each test. (C) UMAP plots of edge and non-edge acinar cells (left) and non-outlier and outlier ductal cells (right). (D) UMAP plots of acinar cells colored by their sample of origin (34 samples in total, as acinar cells from one sample were discarded as they were likely doublets).

We assessed all 9 non-malignant cell types and found that only acinar cells harbored edge cells, having uniquely passed both heterogeneity and proximity tests (Fig. 1B). The existence of edge cells in the acinar population is not due to copy number alterations (CNA) as the acinar cells were shown not to harbor CNAs, in contrast to malignant ductal cells (Fig. S2 in Peng et. al^14^). The non-malignant ductal cells passed the heterogeneity test but not the proximity test, suggesting that ductal cells are highly heterogeneous but that the candidate ductal edge cells do not significantly drift towards malignancy. For clarity, we henceforth refer to the candidate edge ductal cells as outlier ductal cells.

We performed several controls to ensure that the acinar edge population (Fig. 1C) did not arise from common artefacts related to single cell sequencing. Since our analysis is based on acinar cells pooled across non-PDAC and tumor-adjacent PDAC samples, we ascertained that PDAC-adjacent acinar cells are not driving the observed edge-ness in the pooled acinar population (Fig. S2A), nor were edge acinar cells likely to be mis-annotated malignant cells (Fig. S2B). We observed a greater library size and number of transcribed genes in edge acinar cells which may be indicative of their dedifferentiated state^20^. We nevertheless ascertained that the observed differences in library size and expressed gene counts (Fig. S2C) between edge and non-edge acinar populations did not drive the edge behavior. Likewise, four of the non-PDAC samples (samples N1, N2, N3, and N7) had neoplastic indications, which may potentially drive the edge population in pooled samples. However, this was not the case as multiple other samples contributed to the edge cell population (Fig. 1D and Supplementary Section S1). There was no significant difference in the cell cycle status between edge and non-edge acinar cells (Supplementary Section S1), suggesting that cell cycle difference did not drive the edge state. Furthermore, genes known to be up-regulated during tissue dissociation^21^ were not among the genes significantly up-regulated in edge acinar cells.

We note that our computational approach bears similarities to trajectory analysis, where cells whose transcriptomes represent a transition between two cell types can be detected. We thus assessed an analogous trajectory-based pipeline based on Monocle3^22^ for detecting edge cells, where pseudotime values of cells were used to carry out the heterogeneity and proximity tests. This alternative strategy (Supplementary Section S2, Fig. S2D), however, failed to detect edge acinar cells, suggesting that pseudotime values were not suitable for these tests.

Overall, these results reveal an edge subpopulation uniquely in non-malignant acinar cells that have transcriptionally drifted away from the acinar medoid and toward malignant ductal cells. In contrast, ductal cells possess an outlier ductal sub-population that drift away from the ductal medoid but do not drift towards a malignant state.

### Edge acinar cells diverge from a normal acinar phenotype and represent a pre-malignant state

Edge acinar cells expressed PRSS1, a marker of acinar cells, at much lower levels than non-edge acinar cells (Fig. S2B, p < 10^−62^). To further check if edge acinar cells differentially expressed markers of dedifferentiation, we assessed the expression of genes curated by Baldan et. al^23^ that are up- and down-regulated during acinar dedifferentiation. Four genes (*RBPJ,HNF1B,SOX9, MYC*) that are up-regulated during dedifferentiation are also up-regulated in edge acinar cells, while five genes (*AMY2A,RBPJL,SYCN,CPA1,CTRC*) that are down-regulated during dedifferentiation are also down-regulated in edge acinar cells (Fig. 2A). Acinar dedifferentiation precedes acinar-ductal metaplasia --the conversion of acinar to ductal cells during pancreatic injury --which in turn is potentially a precursor to PDAC^24^. We checked expression changes of the genes *STAT3, SEL1L, CBL, KLF4, CTNND1, ICAM1, DCLK1* and *CDKN1A*, which are known to increase in expression during acinar-to-ductal metaplasia^3^. With the exception of *SEL1L*, all other genes were up-regulated in edge-acinar cells (Fig. 2A).

**Fig. 2.**
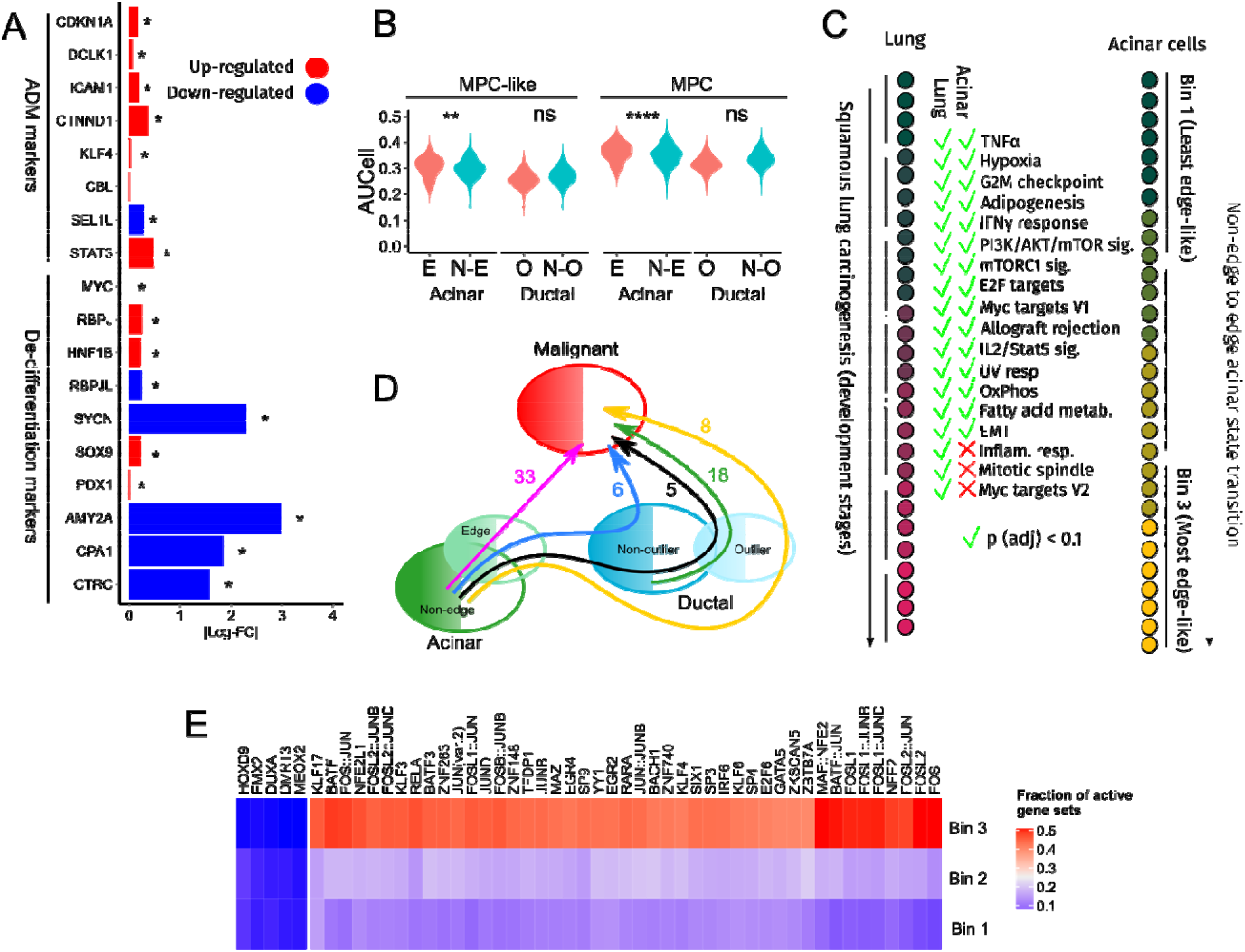
Functional analysis of acinar edge cells. (A) Bars indicate log-fold changes between edge and non-edge acinar cells. (B) The Y-axis is the gene set activity, computed by AUCell, of multi-potent-cell (MPC) and multipotent-cell-like (MPC-like) gene sets across cells in acinar and ductal cell sub-populations shown on the X-axis. **** indicates a p-value less than 10^−4^. (C) Gene sets enriched among genes increasing monotonically in expression during lung cancer progression (Mascaux et. al, left) and from Bin 1 to Bin 3 of the non-edge to edge acinar transition (right). (D) The number of oncogenic gene sets enriched among genes increasing in expression along the cell state transitions indicated by arrows. (E) The fraction of acinar cells in each bin that have an active regulon of the TF indicated along the columns. These are the 50 most variably activated regulons across the three bins.

The acinar cell response during pancreatic injury has been suggested to represent a reversion to a multipotent embryonic progenitor state^25^. Multipotent pancreatic progenitor cells in mice embryos, which differentiate into the main cell types of the pancreas, are marked by expression of Sox9, Ptf1a, Pdx1 and Nkx6-1^26^. We found that SOX9 and PDX1 were up-regulated in edge acinar cells (Fig. 2A), although NKX6-1 showed a negligible up-regulation and PTF1A was down-regulated. Nonetheless, we checked if other genes active in pancreatic progenitors were also expressed in edge acinar cells by processing (see Methods) a single-cell RNA-seq dataset of human fetal (15.4 weeks gestational age) pancreatic tissue^27^. We found a SOX9+PDX1+ cluster (577 cells), which corresponded to cluster #10 in the original publication that the authors termed as multipotent-cell-like (MPC-like) cells. We further refined the MPC-like cluster to require that each cell express all four progenitor markers, yielding 91 *SOX9*+*PTF1A*+*NKX6*-*1*+*PDX1*+ co-expressing cells that we refer to as the multipotent cell (MPC) cluster (Figures S3A,B). We created MPC-like and MPC gene sets from genes up-regulated in the two clusters and scored all acinar cells for activity of both gene sets using AUCell^28^. We found that both gene sets were significantly more active in edge acinar cells than non-edge acinar cells (Fig. 2B). In contrast, among ductal cells, outlier ductal cells did not show higher activity of either of these gene sets compared to non-outlier ductal cells.

Since edge acinar cells are transcriptionally closer to malignant ductal cells than non-edge cells, we checked if the non-edge to edge transition involved known pathways of tumorigenesis. To interpolate intermediate states between non-edge and edge states, we divided acinar cells into three equal-sized bins based on their distance from the acinar cluster medoid. We tested genes monotonically increasing n expression across these bins for enrichment of genes from 50 Hallmark gene sets and 14 gene sets from the CancerSEA^29^ database. Out of 19,276 genes expressed in acinar cells, 3,273 genes exhibited a monotonic increase in expression from the first to the third bin and were enriched for 43 of the 64 gene sets (q-value < 0.1, Supplementary Table 1). We compared our gene expression changes during the non-edge to edge acinar transition with a recent study documenting transcriptomic changes over seven stages of pre-malignancy in the human lung^16^. Interestingly, our findings in acinar cells were highly consistent with those in lung pre-malignant transformation. In Mascaux et. al^16^, the expression of 58% (1848/3366) of genes increased monotonically across the pre-malignant stages, and were enriched for 25 Hallmark gene sets, with 15 gene sets in common with the 43 gene sets in our study (Fig 2C). This included genes related to Myc targets, mTOR signaling, IL2 STAT5 signaling, TNF-alpha signaling via NFKB, response to IFN-gamma, EMT, and UV response.

Next, we investigated four potential paths between non-outlier acinar to malignant cell states (Fig 2D). We identified the genes monotonically increasing in expression along each of these paths and identified enriched oncogenic pathways (Fig. S3C) among these genes. We observed most oncogenic changes (33 pathways enriched) along the path “non-edge acinar -> edge acinar -> malignant” (Fig. 2D). We contrasted this with two other paths, namely, “non-edge -> edge -> outlier ductal -> malignant” and “non-edge -> edge -> all ductal cells -> malignant”, where respectively only 8 and 5 pathways were enriched. This contrast suggests that ductal cells may not always be an intermediate transition state between edge acinar and malignant ductal cells.

We next analyzed transcription factor (TF) activity of acinar cells within each of the three acinar cell bins to understand the transcriptional networks potentially driving the edge acinar state. We processed ATAC-seq data from adult acinar tissue^30^ to first identify 230 TFs whose binding motifs are enriched in ATAC-seq peaks near genes expressed in acinar cells; this analysis also provides putative target gene sets for each TF. We then used AUCell to estimate the fraction of cells in each bin having an active gene set for each of the 230 TFs. We found 50 TFs whose gene sets show the most variable activity among all bins (Fig. 2E, see Supplementary Table 2 for a complete table of all 230 TFs), which can be divided into two groups that are either monotonically increasing or decreasing from Bin 1 to Bin 3. The *RBPJ* gene set showed high activity in Bin 3, which, along with the increase in *RBPJ* expression in edge acinar cells, provides a putative mechanistic link to the re-activation of embryonic progenitor genes in edge acinar cells^31^. The activity of several *KLF* factors increased in Bin 3, including *KLF5*, whose knock-out is known to reduce proliferation in low-grade PanIN cell lines^32^. *HES1* activity, which maintains acinar plasticity^33^, also increased from Bin 1 to Bin 3.

Thus, edge acinar cells differentially up-regulate markers of acinar dedifferentiation and acinar-ductal metaplasia, in addition to the reactivation of genes expressed in embryonic pancreas progenitor cells. This is concomitant with the activation of several oncogenic processes, driven by key TFs, during transition from a non-edge to edge acinar cell state. More surprisingly, there is a substantial commonality between the processes up-regulated in transition from a non-edge to edge acinar cell state and those up-regulated during lung pre-malignant progression.

### Genes up-regulated in edge acinar cells are predictive of PDAC survival

We created gene sets consisting of genes significantly up-regulated and down-regulated in edge acinar cells (Edge-Up and Edge-Down, Supplementary Table 3) and outlier ductal cells (Outlier-Up and Outlier-Down), compared to their respective non-edge and non-outlier counterparts, and analyzed their RNA-seq expression and promoter methylation in both healthy pancreatic tissues and human PDAC tumor samples. We found that Edge-Up genes were up-regulated, while Edge-Down genes were down-regulated in PDAC tumors from the TCGA database, compared to healthy pancreatic tissues from the GTEx database (Fig. 3A). Consistent with gene expression, we found significant hypomethylation at promoters of Edge-Up and hypermethylation of Edge-Down gene promoters in PDAC samples (Fig. 3B). This suggests that gene expression and methylation changes in acinar edge cells foreshadow changes in PDAC tumors in a consistent manner.

**Fig. 3.**
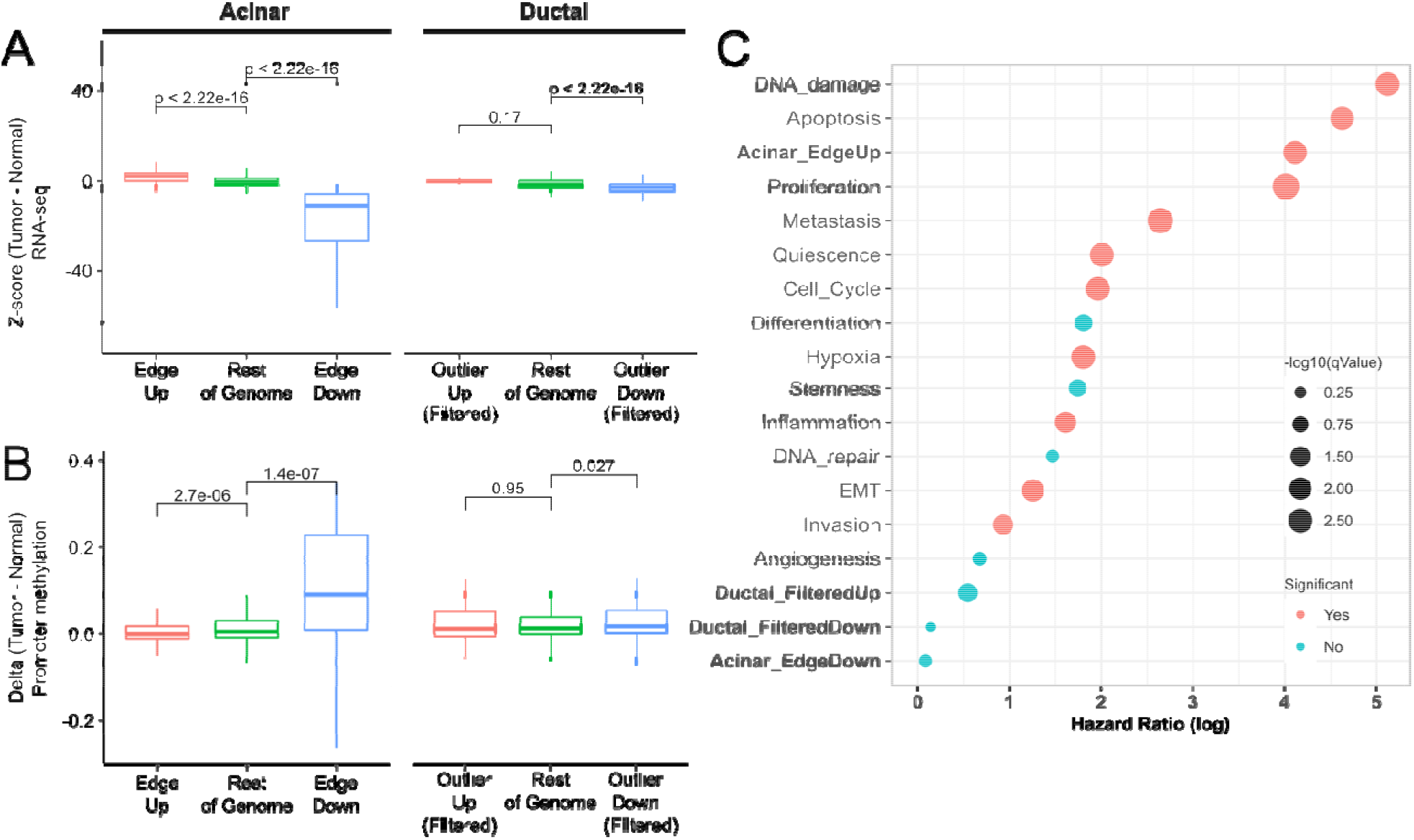
Acinar Edge and Ductal Outlier genes in TCGA PDAC. (A) RNA-seq expression z-scores in PDAC samples (using GTEx pancreas RNA-seq as a reference) of up-regulated (red), down-regulated (blue) and remaining (green) genes in edge-acinar cells and outlier-ductal cells. Genes in the Outlier-Up and Outlier-Down datasets are filtered to remove overlapping Edge-Up and Edge-Down genes. (B) Methylation z-scores among PDAC using methylation samples from healthy samples as a reference (see Methods) of gene promoters in A, (C) Log of Hazard ratios obtained from Cox regression of gene sets in TCGA PDAC samples.

When we repeat these analyses for ductal Outlier-Up and Outlier-Down gene sets, counterintuitively (since outlier ductal cells do not exhibit a drift towards malignancy), we found a similar trend as for acinar cells, where Outlier-Up genes were up-regulated while Outlier-Down genes were down-regulated in PDAC tumors (Fig. S3E), and Outlier-Up gene promoters were hypomethylated (Fig. 3B), though Outlier-Down gene promoters were not hypermethylated. We scrutinized these counter-intuitive observations and found that this is likely because over half the Outlier-Up genes were also Edge-Up genes, with only 8 Outlier-Up (and 177 Outlier-Down genes) being ductal-specific in their expression pattern. Removal of these overlapping genes eliminates these trends in RNA-seq and methylation patterns (Fig. 3A,B).

We further assessed, using a Cox proportional-hazards model, whether the four gene sets’ activity in PDAC tumors are associated with patient survival. As shown in Fig. 3C and Fig. S3E, both Edge-Up and Outlier-Up gene sets have a significant hazard ratio (q-value < 0.1), but Edge-Up gene set has a higher hazard ratio than Outlier-Up genes. Notably, neither Edge-Down nor Outlier-Down gene sets are significantly associated with survival. As above, repeating the survival analysis based on ductal-specific Outlier-Up genes does not show significant association with survival. We also performed Cox regression for oncogenic gene sets in CancerSEA and found that a majority of these sets were predictive of survival, albeit with a lower hazard ratio than the Edge-Up gene set.

These results suggest that the genes increasing in expression in the edge-acinar state were key to tumor progression and are in line with our findings (Fig. 2C) that several oncogenic processes are enriched only among genes increasing in expression during the non-edge to edge transformation.

### Acinar edge cells are found in independent healthy pancreas samples

We checked if edge states can be found among acinar cells in other published single-cell datasets of human pancreatic tissues. We re-analyzed published SMART-seq^26^ (GSE81547) and CEL-seq^34^ (GSE85241) single-cell RNA-seq datasets of healthy human pancreas samples. To check for the presence of edge cells, we used AUCell to score acinar cells in both datasets based on our signature gene set. We then declared each cell as edge or non-edge based on the Global_k1 activity threshold computed by AUCell. First, similar to Fig. 2A, we compared the log-fold changes of acinar-ductal metaplasia and acinar dedifferentiation markers between edge and non-edge acinar cells (Fig. 4A). In GSE81547, all 9 dedifferentiation markers, and 6 out of 9 ADM markers, showed consistent fold-changes with edge acinar cells. In GSE85241, 6 out of 9 dedifferentiation markers, and 4 out of 9 ADM markers, showed consistent fold-changes with edge acinar cells.

**Fig. 4.**
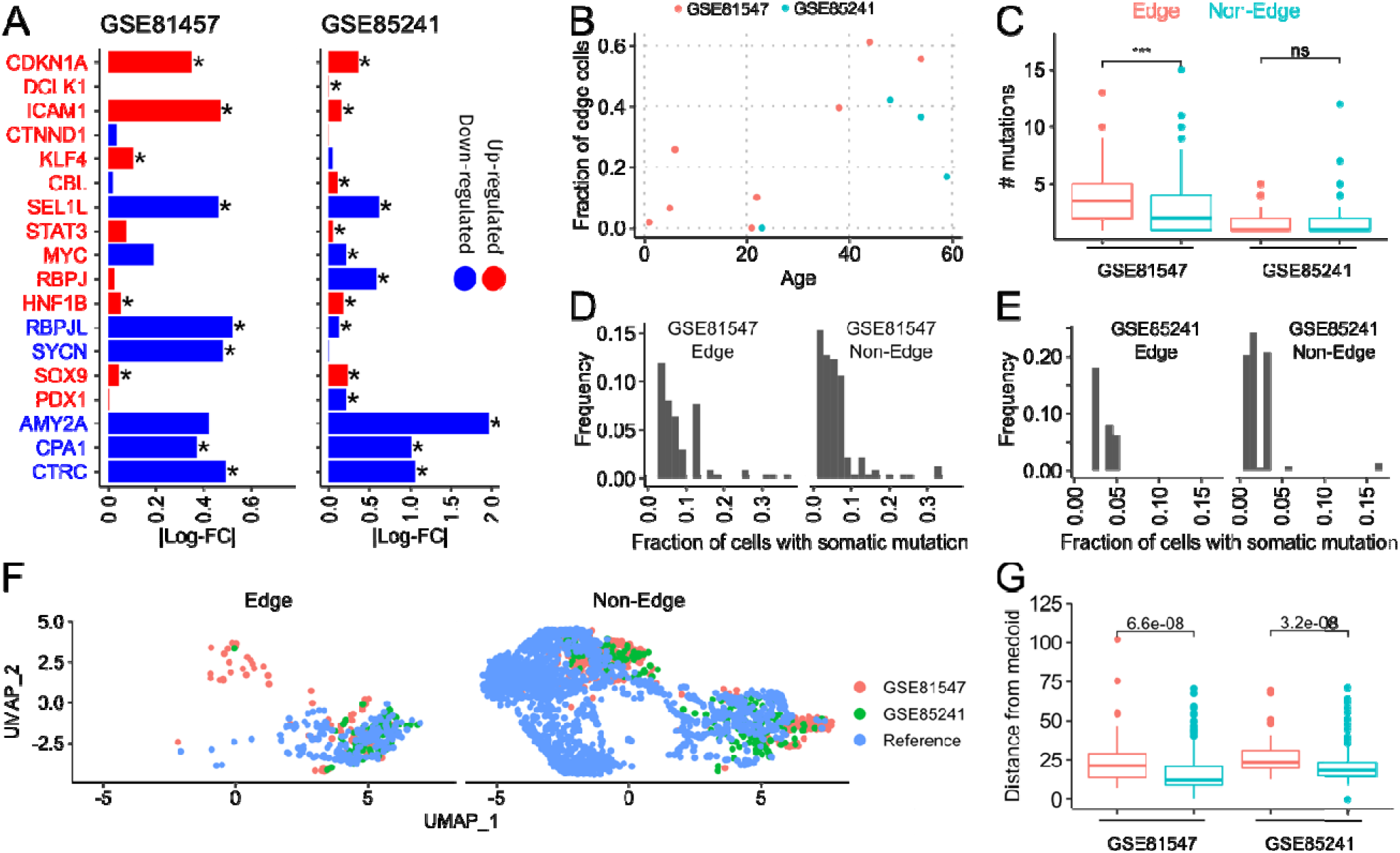
Acinar edge cells in independent datasets and links with aging. (A) Bars indicate log-fold changes between edge and non-edge acinar cells in GSE81547 and GSE85241. Markers in red and blue fonts are known to be up-regulated and down-regulated, respectively, during ADM (CDKN1A to STAT3) and dedifferentiation (MYC to CTRC). Matching color of the marker text and the bar indicates that the observed log-fold change matches the expected gene expression change of the marker. (B) Scatter plot of fraction of edge-acinar cells with the age of the tissue donor in GSE81547 and GSE85241. (C) Number of mutations in edge and non-edge acinar cells in GSE81547 and GSE8524. (D,E) Histogram of the fraction of edge cells and non-edge cells that contain a somatic mutation. (F) UMAP visualization of acinar cells from GSE81547, GSE85241 and the reference dataset from which the edge signature was derived. (G) Distance of edge (red) and non-edge (cyan) cells from the medoid acinar cell in PCA space computed separately for GSE81547 and GSE85241 datasets.

Since PDAC risk increases with age, we checked if there was an age-dependent increase in the fraction of edge cells in these datasets. Since our edge signature is derived from patients spanning five decades of age, to remove any age-associated confounding factor from our edge signature, we carried out a linear regression of gene expression against age (along with the number of expressed genes and cell cycle scores as covariates). This removed 8 genes from the acinar edge signature that were increasing in expression with age (FDR < 0.1). When we used AUCell to determine edge cells in both GSE81547 and GSE85241 datasets using this refined signature, we found a significant age-dependent increase in the fraction of edge cells (R^2^ = 0.66, p = 0.02) across both datasets (Fig. 4B).

Since tissues also accumulate somatic mutations during aging, we assessed whether the edge cell state may be driven by somatic mutations, i.e., whether edge cells harbor differential somatic mutations. We called variants in acinar cells using the GATK Best Practices pipeline (Methods). The number of somatic mutations in these cells agreed with estimates of somatic mutations rates in pancreas tissue in GTEx data^35^. We found that edge acinar cells had more somatic mutations than non-edge acinar cells in GSE81547 but not in GSE85241 (Fig. 4C). The differences between both datasets likely stem from differences in their library sizes, with GSE81547 being sequenced to a much higher depth^36^. Nonetheless, in both datasets, all these mutations were rare, and were present, on average, in 2.18% and 3.47% of non-edge and edge cells in GSE85241, and in 6.34% and 8.53% of edge cells, respectively in GSE81547 (Figures 4D,E), which does not support a clonal origin for edge cells. This modest difference in mutation frequency between edge and non-edge cells is not significant based on sampling that preserves the number of edge and non-edge cells in each sample. Further, none of the mutations in edge and non-edge cells were classified as oncogenic driver mutations in the COSMIC cancer gene census (v92)^37^.

We compared the edge cells from these two datasets with the edge cells found in our reference dataset. We used Seurat to integrate acinar cells across all three datasets and found that the edge cells (and non-edge cells) overlapped each other in UMAP space (Fig. 4F). Further, within both GSE81547 and GSE85241 acinar cells, the edge cells were significantly farther from their respective medoids than non-edge cells (Fig. 4G). Thus, the edge states in each of these datasets are similar and represent a transcriptional drift away from the normal acinar state in each of them.

These findings validate the existence of edge-like acinar subpopulation cells in additional datasets, where they consistently exhibit expression profiles of ADM and dedifferentiation markers as in the PDAC dataset. Furthermore, we observe a strong correlation between frequency of edge cells and age.

### Edge-like variation in non-malignant liver and lung tissues

Alveolar type 2 (AT2) cells are believed to be the cell-of-origin^38^ of lung adenocarcinoma (LUAD) tumors. However, application of our original pipeline on scRNA-seq data from non-malignant (AT2) and LUAD samples^39^ did not detect an edge sub-population among AT2 cells, or any other non-malignant lung epithelial cluster. We then modified our original pipeline to check if any individual principal components reflected significant gene expression heterogeneity and a drift towards malignancy. Here, heterogeneity and the proximity tests are done for individual Normal and Pooled PCs respectively, and an additional test of collinearity between the qualifying Normal PC and the qualifying Pooled PCs (Fig. 5A, Methods). We note that multiple Normal PCs can show heterogeneity and drifts towards malignancy, reflecting the activation and inhibition of different gene sets in a subset of non-malignant cells. With this refined pipeline, we found that Normal PC5 of AT2 cells defined an edge population that showed a drift towards a malignant cell cluster (Tumor State 2) along Pooled PC 1, which is collinear with Normal PC5 (Correlation coefficient = 0.82, q-value < 10^−9^). Additionally, Normal PC1 of Club cells, and Normal PCs 1 and 2 of AT1 cells, also represented drifts towards malignancy.

**Fig. 5.**
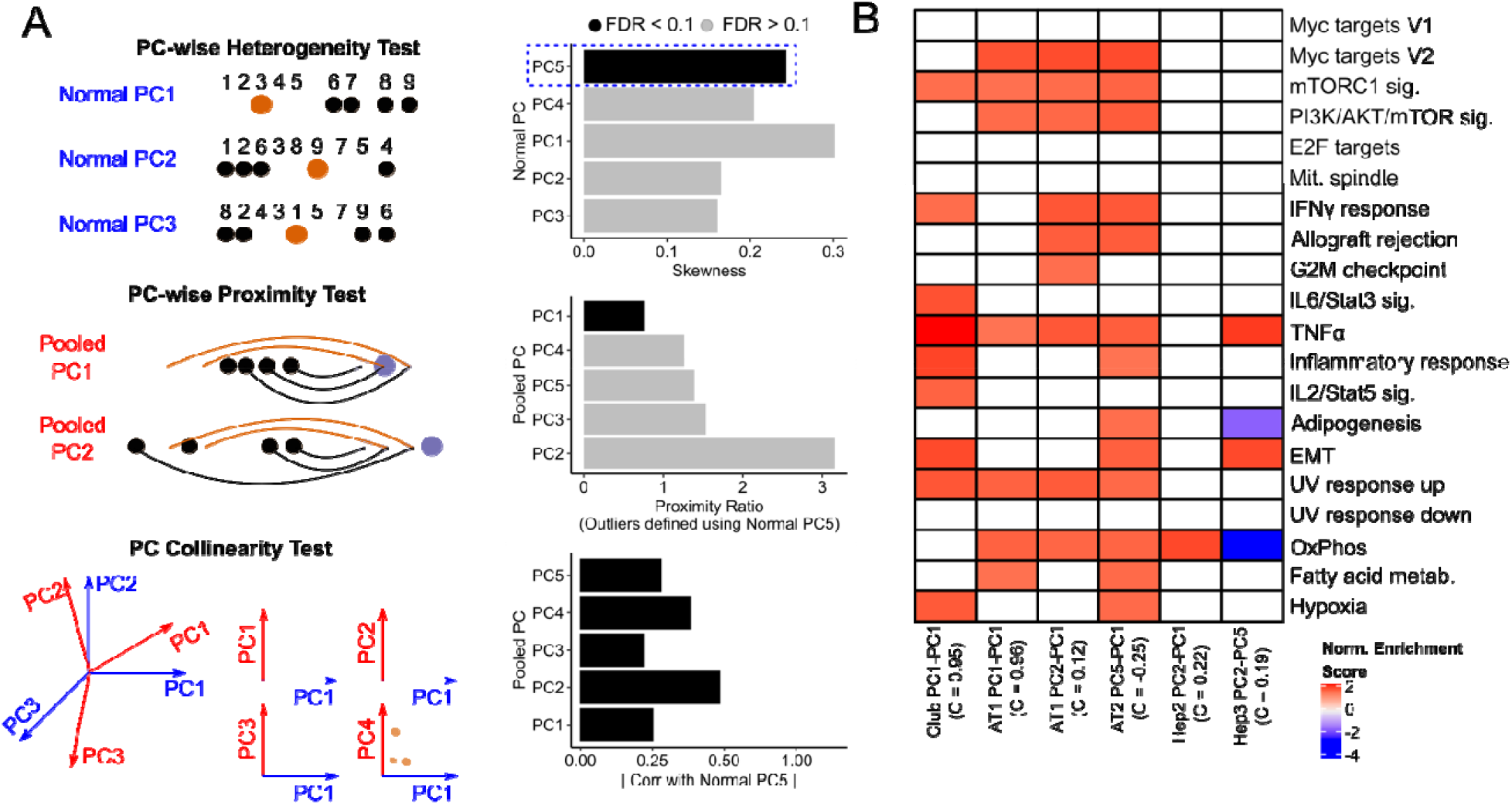
Edge heterogeneity in lung epithelial cells and hepatocytes. (A) Schematic of three-stage pipeline to detect directions of edge heterogeneity. Each Normal PC is tested for positive skewness, and each PC that passes the test is used to define an outlier cell population (in black). For each outlier population, each pooled PC is then used to compute distances between non-malignant and malignant cells and carry out the proximity test, with all PCs tested for collinearity with the Normal PC used to define the outlier cells. Collinearity is defined as the Spearman correlation between the Normal and Pooled PC scores of all non-malignant cells in the cluster. Those skewed Normal PCs that are collinear (FDR < 0.1) with a Pooled PC that passe the proximity test represent directions of edge heterogeneity within the non-malignant cluster. The bar plots shown are from running the three-stage test on Alveolar Type 2 (AT2) cells, where Normal PC 5 is used to define outlier cells (B) Normalized enrichment scores of gene sets (those that are active during lung cancer progression in Mascaux et. al) along Normal PCs passing all three tests amongst Club, AT1 and AT2 cells in lung, and Hep2 and Hep3 cells in liver. The Normal and Pooled PC pair that pass the heterogeneity tests and proximity tests are indicated, along with the collinearity score between the two qualifying PCs. The collinearity scores all have a q-value < 0.1.

Next, we tested non-malignant liver epithelial cells for an edge-subpopulation that show a drift towards liver hepatocellular carcinoma (LIHC). We pooled single-cell RNA-seq data from healthy liver^40^ and LIHC samples^41^ to create a dataset with both healthy and malignant cells (caveats with this dataset are discussed in Supplementary Section S3). We found that Normal PC2 of two hepatocyte clusters, Hep2 and Hep3, exhibited drifts towards malignancy along Pooled PC1 and PC4, respectively.

We used each of these Normal PCs that exhibited a drift toward malignancy to define an “edge-like” population as before and tested the genes up-regulated in the edge-like cells for enrichment of Hallmark and CancerSEA gene sets (Supplementary Figure S3D). We found that edge-like AT2 cells (based on Normal PC5) and AT1 cells (based on Normal PC2) were enriched (Fisher test, p = 0.042 and p=0.05, respectively) for gene sets active in lung cancer progression in the Mascaux et. al study (Fig 5B). This was however not the case for edge cells identified in Hep2 and Hep3 cell types.

Overall, while we did not find prominent edge cells in lung and liver based on global transcriptomic shifts, our results nevertheless suggest significant heterogeneity in specific oncogenic programs in non-malignant epithelial cells of lung and liver.

## Discussion

Here we show the existence of a subset of non-malignant acinar cells that we refer to as edge cells^15^, that are transcriptionally distinct from a typical acinar cell, and significantly closer to malignant PDAC cells. This phenomenon is observed broadly across individuals and in multiple datasets. Although edge cells do not seem to be driven by clonal somatic mutations, interestingly, we see evidence of increased prevalence of edge cells with age, and consistently, an enrichment of edge-up-regulated genes among genes increasing in expression with age.

One way to interpret the observed global transcriptional drift in acinar edge cells toward malignancy is that there are overlapping oncogenic programs that individually show heterogeneity in the non-malignant cell population and are broadly concordant with each other. Ultimately, an increased transcriptional activity along multiple oncogenic programs in a subset of cells is revealed as the edge cells by our approach. Our results also reveal significant heterogeneity involving several oncogenic programs in non-malignant epithelial cells of lung and liver.

There is a significant overlap between pathways activated in the non-edge to edge transitions in acinar cells on the one hand, and those activated during pre-malignant progression in the lung on the other. In acinar edge cells, we see an up-regulation of the targets of transcription factors *RBPJ, HES1*, and *KLF5* targets, which are known to mediate acinar cell plasticity^33^ and a reversion to a multi-potent pancreatic progenitor state^31^. This suggests a role for known transcriptional networks playing a role in the transition to an edge state. It has been proposed, and experimentally shown, that transcriptional fluctuations within complex regulatory networks can result in a clonal population existing in multiple phenotypic states^9,10,42^. In cancer cells, such fluctuations can cause cells to switch between drug-sensitive and drug-resistant states^43^ in a manner that can be perturbed by targeting key transcription factors^44^.

Once a cell transitions to an edge state, how long a cell spends in an edge state, and whether the edge state persists after cell division, may additionally involve epigenetic mechanisms. DNA methylation and epigenetic inheritance through histone modifications represents one such potent mechanism^45,46^. Coupled single cell transcriptomics and DNA methylation data from the same acinar cell, which is needed to precisely assess the role of DNA methylation in sustaining the edge cell population, is currently not available. However, we found that the promoters of genes that are up-regulated in the edge acinar cells relative to non-edge cells, were hypomethylated in PDAC tumors, and the converse was true for genes down-regulated in edge acinar cells, suggesting a potential role of epigenetics in maintaining the edge cell state.

A potential role of the tissue environment, and DNA methylation, in giving rise to edge cells is further supported by our observed link between age and the fraction of edge cells in healthy acinar cells. Aging is the greatest risk factor for most cancers^47^. While clonal expansion of somatic mutations does occur with age in certain tissues such as skin and oesophagus^48^, we found no evidence of clonal expansion in the edge acinar cells. Beyond the role of mutations, epigenetic changes from age-related hypomethylation^49^ likely contribute to the stability and rate of switching to an edge state with age.

While we have demonstrated, in multiple contexts, that a specific edge cell subpopulation in healthy tissues is transcriptionally distinct and is significantly closer to malignant cells, it is not clear if, and precisely how, the edge cells may contribute to tumorigenesis. Many processes that robustly change with age, such as genomic stability, telomere attrition, metabolism, and cellular senescence, are also mechanistically linked to cancer^17^. One possibility is that the edge cells are in an epigenomic state that is more sensitive to oncogenic mutations. For instance, cigarette smoke alters DNA methylation patterns in epithelial cells and sensitizes them towards KRAS-induced transformation^13^. Thus, even though the emergence of the edge transcriptional state may not be driven by mutations, the edge state may be more sensitive to oncogenic transformation with age-associated transcriptomic changes being more sensitive to such mutations^50,51^.

Overall, our results support the notion of an edge transcriptomic state in healthy tissues that is pre-malignant. Pancreatic acinar cells likely switch between edge and non-edge states, although the time spent by cells in either state is unclear. Establishing the stability of these states would require the tracing of lineages of acinar cells to infer the regulatory changes underlying the switching process.

## Acknowledgments

This work utilized the computational resources of the NIH HPC Biowulf cluster and was supported by the Intramural Research Program of the National Cancer Institute, Center for Cancer Research, NIH. We would like to thank Curtis Harris, Xin Wang, Shouhui Yang and Cenk Sahinalp for discussions and feedback. We especially thank Argiris Efstratiadis for his feedback and comments on our re-analysis of fetal pancreatic single-cell RNA-seq data. We thank Arati Rajeevan for help with illustration.

## Methods

The code necessary for reproducing these results are available at https://github.com/hannenhalli-lab/pdac_edge.

### Processing PDAC scRNA-seq data (CRA011600)

We downloaded processed single-cell RNA-seq read counts and cell type annotations from (ftp://download.big.ac.cn/gsa/CRA001160). Out of 57,730 cells, we removed 2,877 cells that were declared as doublets with doubletFinder v3.0 (assuming a prior 5% doublet rate). In all downstream calculations, we used the default Seurat (v3.0) normalization scheme implemented in FindNormalize (with a scale factor of 10,000) to normalize library size variation across cells.

### Two-stage statistical test for an edge sub-population

Our procedure for testing whether a non-malignant cell cluster harbors an edge sub-population consisted of two tests ---the skewness and the proximity tests.

#### Heterogeneity test

We selected the 1000 most variable genes (using Seurat’s default FindVariableFeatures function) in the non-malignant cell cluster, z-score normalized their expression, and computed a 50-dimensional PC embedding for each cell; we refer to these PCs as Normal PCs to underscore that they are computed only from the non-malignant cell cluster. We then computed the distance of each cell from the cluster medoid based on Euclidean distance. The 10% of cells that are farthest from the medoid are termed outlier cells. We quantified heterogeneity as the skewness, *s*, of the distance distribution using the medcouple estimator from the robustbase package in R. To compute the statistical significance of *s,* we create 100 control cell clusters by shuffling each of the 50 Normal PC coordinates across all cells in the original cluster. For each control cluster, we compute the skewness as above, and based on a Gaussian fit of these 100 control skewness values, we estimated the empirical p-value of *s*. We used a p-value threshold of 0.01 to consider the cell cluster heterogeneous and proceed to the next test.

#### Proximity test

Here we determine whether the outlier cells in the non-malignant cluster are significantly closer to the malignant cell cluster than non-outlier cells. We carry out PCA jointly on both malignant cells and non-malignant cells, using 1000 most highly variable genes across these cells. We refer to these PCs as Pooled PCs. We then define the malignant cell cluster’s medoid using the Euclidean distance metric, and compute the proximity ratio, R, as the ratio between the average distances of outlier cells (in the Pooled PC space) to the malignant cluster medoid to that of the non-outlier cells. A value of R < 1 implies that the outlier cells are closer to malignancy than non-outlier cells. We compute the statistical significance of R by randomly choosing 10% of the non-malignant cells as outlier cells, and re-compute R using these control outlier cells. We repeat this process 100 times, fit a Gaussian to the obtained ratios and estimate the empirical p-value of observing a value less than R. If this p-value is less than 0.01, the outliers are labelled as edge cells.

### Modified three-stage statistical test for finding edge heterogeneity

The three-stage pipeline retains the heterogeneity and proximity tests and incorporates a third collinearity test.

#### Heterogeneity test

We computed 5 Normal PCs based on the 1000 most variably expressed genes within the non-malignant cluster of interest. Using each PC individually, as above, we defined the medoid cell, computed the distance of each cell from the medoid, followed by skewness of the distance distribution, s, and its significance based on shuffling the expression separately amongst cells in each sample (this sample-aware shuffling removes any potential bias caused by inter-sample heterogeneity). The p-values of *s* computed for all 5 PCs are corrected using the Benjamini-Hochberg FDR procedure. Each Normal PC with an FDR < 0.1 is chosen to define outlier cells, i.e., 10% of cells farthest from the cluster medoid.

#### Proximity test

5 Pooled PCs are computed based on the 1000 most variably expressed genes across the pooled non-malignant and malignant clusters. For each outlier cell population (defined by a particular Normal PC qualifying the Heterogeneity test), the proximity ratio of the outlier cells, R, and its p-value, is computed separately for each pooled PC as above. The FDR is then computed for each pooled PC, and the set *P* of all Pooled PCs with an FDR < 0.1 are retained.

#### Collinearity test

We compute a 5×5 correlation matrix of Spearman correlation coefficient between every Normal and Pooled PC score pair across all cells in the non-malignant cluster that qualify both heterogeneity and proximity tests. The p-values of each correlation is corrected using the FDR method. For each skewed Normal PC, if there exists at least one collinear Pooled PC with a low proximity ratio (with a correlation FDR < 0.1), then the Normal PC is a direction of edge heterogeneity.

### Processing fetal pancreas data

We downloaded the cell-by-gene read count matrix from the GEO database (accession number GSE141087). We first removed cells that had more than 10% of their reads aligned to mitochondrial genes, after which we removed doublets using DoubletFinder v3.0 (50 principal components, 5% doublet rate and p_N = 0.25). We normalized the read counts using Seurat’s default FindNormalize function, after which we selected 1000 most highly variable genes and carried out PCA with 50 PCs. We then clustered the cells using the FindClusters function in Seurat (resolution = 0.8, k = 30 neighbors in PC space for constructing the neighborhood graph). We then scored each cluster based on the average normalized expression of markers of multi-potent progenitors (NKX6-1+SOX9+PTF1A+PDX1+) known from mouse experiments, yielding 577 multipotent-like (MPC-like) cells, of which 91 cells co-expressed all four marker genes. We then used the FindMarkers function from Seurat to find genes that were up-regulated in these two progenitor clusters which were then used as gene sets to score all acinar and ductal cells using AUCell.

### Gene set enrichment comparison to Mascaux et. al

We divided the acinar cells into three bins based on their distances from the acinar medoid in PC space. We z-scored the normalized expression of each gene across all acinar cells and picked genes that increased in z-score by at least 0.1 between adjacent bins. We carried out a Fisher test for over-representation for 64 gene sets (50 Hallmark gene sets and 14 CancerSEA gene sets), after which we carried out an FDR correction and picked gene sets with a q-value < 0.1 as significant.

### Motif enrichment and network analysis

We downloaded acinar-specific ATAC-seq reads from GEO (<u>GSM1606431</u>). Reads were trimmed with trimgalore (v0.6.5) and aligned to the human genome (hg19) using BWA^52^ (v0.7.17). MACS2^53^ (v2.2.6) with default parameter settings was used to call peaks with a q-value < 0.05

To find a list of motifs enriched near acinar-expressed genes, we used the SPRY-SARUS motif scanner^54^ to scan the central 100 bp region of ATAC-seq peaks for matches to motifs in the JASPAR 2020 vertebrate motif collection^55^. Out of 746 motifs, we restricted our scans to 589 motifs that involved a TF that was expressed in at least 10% of all acinar cells. We split the ATAC-seq peak regions into foreground or background sets depending on whether or not the peaks were at most 10kb upstream of a gene expressed in at least one acinar cell. We scanned both sets of regions for motif matches (p-value < 10^−4^) and carried out a Fisher test of over-representation among the foreground sequences for each motif. We then computed q-values for each TF and retained TFs with a q-value < 0.1.

For each retained enriched TF, we created gene sets that consisted of its putative gene targets in the foreground set. We scored each gene set’s activity in each acinar cell using AUCell and used AUCell’s internal Global_k1 threshold to declare a gene set as active or inactive in each acinar cell. We then computed the fraction of acinar cells in each of the 3 bins with an active gene set, with the same cell -bin assignment that was computed in Fig. 2C.

### Processing aging pancreas single-cell RNA-seq data

We downloaded processed single-cell RNA-seq read counts from GEO GSE81547 and GSE85241. For GSE81457, we used prior annotations of cell types from the original publication^26^ to find transcriptomes of acinar cells. For GSE85241, we clustered cells using Seurat with the same parameters as we used for the fetal pancreas scRNA-seq data. As in the original publication, we picked the cluster with the highest PRSS1 expression as the acinar cell cluster.

### Variant calling

We called variants in acinar cells from the raw sequencing reads in GSE81547 and GSE85241 datasets using the GATK best practices workflow. We then removed variants that were (a) shared across donors, (b) were annotated in dbSNP v138, (c) had fewer than 5 reads aligning to the locus or had fewer than 3 reads supporting the alternate allele.

### Processing lung and liver scRNA-seq data

Lung: For the lung adenocarcinoma set, we downloaded processed read counts and cell type annotations from GSE131907. The malignant cells were annotated as tS1, tS2 or tS3 (Tumor States 1,2 and 3). We chose tS2 cells as the malignant reference since they represented a more transformed malignant state^39^.

Liver: Healthy liver data was downloaded from https://github.com/BaderLab/HumanLiver in the form of a Seurat object that contained cell type annotations and read counts for each cell. Processed hepatocellular carcinoma read counts and annotations were downloaded from GSE125449.

### Differential expression analyses between edge and non-edge cells

In acinar and ductal cell populations, genes differentially expressed respectively in edge and outlier cells were found using the “LR” test in the Seurat FindMarkers function, where the p-value associated with each log-fold change was estimated after controlling for cell cycle scores, number of expressed genes and inter-sample variation between cells.

In AT2 and Hep2 cells, we found the LR and Wilcoxon tests in the Seurat FindMarkers function to be overly conservative in computing the significance of log-fold changes. Hence, for these two cell types, we employed MAST^56^ to find differentially expressed genes between edge and non-edge cells after controlling for inter-sample variation between cells (effects of cell cycle and number of expressed genes was negligible and thus not modelled).

### Survival analysis of TCGA cancer patients

For each cancer type investigated here, we obtained the mRNA expression (in TPM units) and clinical data for TCGA cancer patients from UCSC-xena browser *(https://xena.ucsc.edu/)*. We used Cox regression to model the overall survival of patients by using the median expression of each signature gene set (y-axis of Fig. 3C) as an explanatory variable. Additionally, we used the age of patients as covariate and stratified the model based on their gender to control for these potential confounders. The resulting p-values were corrected for multiple comparisons using the FDR method and hazard ratios were plotted on log scale.

### Expression analysis in bulk tumor data

For each cancer type investigated, we z-scored the expression of each gene in TCGA cancer patients based on its mean and standard deviation in normal samples of corresponding tissue from Gtex and used the averaged z-scores to compare different gene sets. Prior to z-scoring, we performed quantile normalization in order to make the two datasets comparable.

### DNA methylation analysis in bulk tumor data

We used 450k DNA methylation data of cancer and normal samples from array-expression for pancreas^57^ and from GEO database for lung (GSE66836) and liver (GSE54503) samples. The coordinates of 450k methylation array probes were obtained using the COHCAP library in **R** and were mapped to the 5kb upstream promoter region of each gene using bedtools. We used the mean and standard deviation of aggregated methylation of each promoter in normal samples to compute the z-scores of the same in the cancer samples and plotted the averaged z-scores to compare different gene sets.

