## Supplement for "A transcriptionally distinct subpopulation of healthy acinar cells exhibit features of pancreatic progenitors and PDAC"

### **Supplementary Notes**

#### **S1. Tests for potential confounders that may give rise to an edge population**

We performed several controls to ensure that the acinar edge population did not arise from any artefacts related to single cell sequencing.

***Tumor-adjacent cells not likely to drive edgeness in acinar cells.*** Our non-malignant acinar populations comprised both non-PDAC samples as well as tumor-adjacent samples from PDAC patients, raising the concern that tumor-adjacent acinar cells may be driving the edge-ness. We assessed this possibility through following analyses. First, we found that acinar cells still had an edge sub-population even after all tumor-adjacent (TA) acinar cells were excluded (Skewness = 0.5313 (p = 5.44 x 10^-6^), Malignant-proximity ratio = 0.9611 (p = 3.319 x 10^-9^). Second, the non-edge to edge differential expression across genes is highly concordant with and without the TA cells (Log fold R^2^ = 0.93, p < 10^-10^, Fig. S2A). Finally, TA-acinar cells were significantly closer to the medoid of the acinar cluster than the medoid of the malignant cluster (Fig. S2B, right column). Together, these three analyses suggested that TA-acinar cells are transcriptionally more similar to non-edge acinar cells and their inclusion did not erroneously give rise to an edge sub-population.

***Acinar edge cells not likely to be mis-annotated malignant ductal cells.*** In the original publication^14^, acinar cells were annotated based on expression of the marker genes PRSS1, CTRB1, CTRB2 and REG1B, while malignant ductal cells expressed the marker genes KRT7, KRT19, TSPAN8 and SLPI, and harbored copy-number variations. We found that the acinar edge cells expressed the acinar marker genes (Fig. S2C, bottom row) at a significantly lower level than non-edge cells and at a significantly higher level than malignant ductal cells. Further, edge acinar cells expressed malignant ductal markers at a significantly higher level than non-edge cells and at a significantly lower level than malignant cells. Along with the observation that the acinar cells still harbored an edge sub-population after the exclusion of tumor-adjacent cells, this suggested that the edge acinar cells were not actually malignant cells that were mis-annotated as acinar cells.

***Library size and expressed gene counts not likely to drive edge-ness*.** We observed that edge acinar cells have a greater library size (number of mapped sequencing reads) than non-edge acinar cells (Edge : 8035 (25th percentile) -- 24238 (75th percentile), Non-Edge : 4625 (25th percentile) -- 11874 (75th percentile)). Edge acinar cells also had a higher number of expressed genes (2182 (25th percentile) -- 4456 (75th percentile)), than non-edge cells (792 (25th percentile) -- 1927 (75th percentile)). A higher gene count is consistent with a dedifferentiated or stem-like state^20^. However, to ascertain that the edge cells were not simply found as edge because of their greater library size or expressed gene counts, we separately under-sampled reads and expressed genes in the edge acinar cells such that their library sizes and expressed gene counts were identically distributed to those of non-edge cells (Fig. S2C). We then re-ran the edge statistical tests on the acinar cell cluster, using the resampled cells in place of original cells, and computed the p-values of the skewness and malignant-proximity ratio tests. We repeated this re-sampling process 100 times and computed q-values for both tests. We found these p-values to be less than 0.1 in all 100 replicates, suggesting that read count and expressed gene count differences were not artifactually giving rise to an edge population.

***Cell cycle heterogeneity not likely to drive edge-ness****.* We checked if there were significant differences in proportions of cells in G1, G2/M and S phases between edge and non-edge acinar cells. We used the CellCycleScoring function in Seurat to classify all acinar cells as being in either G1,G2/M or S phases. There were 618, 548 and 499 non-edge cells in G1, G2/M and S phases, respectively, while the corresponding numbers for edge cells were 82,72 and 30. We carried out a binomial test between the fractions of cells in each of these phases between edge and non-edge cells and found no significant differences in the proportions (p = 0.36 for cells in G1, p=0.33 for cells in G2/M, and p=0.30 for cells in S phase).

***Batch/inter-sample effects not likely to drive edge-ness.*** Since the 1,841 acinar cells used in our analyses were derived from 35 samples, we checked if edge acinar cells were arising from multiple samples or from a single sample. We expected that the fraction of edge cells arising from each sample would be proportional to the fraction of acinar cells arising from that sample. We used this null expectation to simulate 10000 draws of 184 edge cells from all 35 samples, drawn according to proportions of acinar cells coming from each sample. For each draw, we computed the entropy of the distribution of the fraction of edge cells from each sample. From this distribution of entropies (Fig. S2E, approximated by a Gaussian distribution, mean = 2.23 bits, deviation = 0.082 bits), the left-tailed p-value of the observed entropy of the fraction of edge cells was 0.504, suggesting that the distribution of edge cells among different samples was consistent with chance.

#### **S2. Two-stage statistical test based on Monocle3 trajectory analysis**

We checked if pseudotime values assigned to acinar and ductal cells using the Monocle3 trajectory analysis too^22^ could be employed to find edge acinar cells. Below, we describe versions of the heterogeneity and proximity tests based on pseudotime.

For the heterogeneity test, analogous to using Normal PCs, we constructed a trajectory of only non-malignant cells (either acinar or ductal cells), with the cell closest to the medoid of the non-malignant cell cluster being assigned as the root cell i.e., a pseudotime value of 0. We then computed the skewness of the pseudotime distribution and tested it for a significantly positive skew. The control skewness distribution was constructed by separately computing the skewness of 100 trajectories of non-malignant cells with cross-cell shuffled principal component projections. If the p-value of the observed skewness was less than 0.05, we considered the non-malignant cells to be significantly heterogeneous, and the 10% of cells with the highest pseudotime were labelled as outlier cells.

For the proximity test, again, analogous to our standard pipeline, we constructed a trajectory of both non-malignant and malignant cells, with the cell closest to the medoid of the non-malignant cell cluster assigned as the root cell. We then computed the proximity ratio as the ratio between the mean squared difference between all outlier cells’ pseudotimes and the median malignant cell pseudotime to those of all non-outlier cells. We tested if the ratio was significantly low by computing the proximity ratio of outlier and non-outlier cells after randomly choosing 10% of non-malignant cells as outlier cells. If the p-value of the proximity test was less than 0.05, we considered the outlier cells to be an edge cell population.

All trajectories were constructed using 50 principal components computed from log-normalized read counts. The align_cds function was used to correct for any batch effects and regress any residual cell cycle differences that may influence the pseudotime.

When we tested both acinar and ductal cells for edge sub-populations using these two tests, we found that neither acinar nor ductal cells passed the heterogeneity or proximity tests (Fig. S2D).

#### **S3. Untested confounders in liver scRNA-seq analysis**

There are two related caveats to results from the human liver. First, the scRNA-seq data used for non-malignant liver cells possibly contains uncorrected batch effects, since all Hep2 cells were found to come from a single donor, while other hepatocyte clusters came from multiple donors. Second, the scRNA-seq data that contained malignant cells did not include any hepatocytes. Since batch correction requires the same cell types to be present across all donors, we could not apply any batch correction technique. This means that the Pooled PCs used to carry out the proximity test are likely to represent, in part, batch variation and not just variation between non-malignant and malignant cells. In contrast, this was not an issue with our PDAC and LUAD analyses, since tumor biopsies included sufficient adjacent-normal epithelial cells to overcome any batch-related variation in the Pooled PCs.

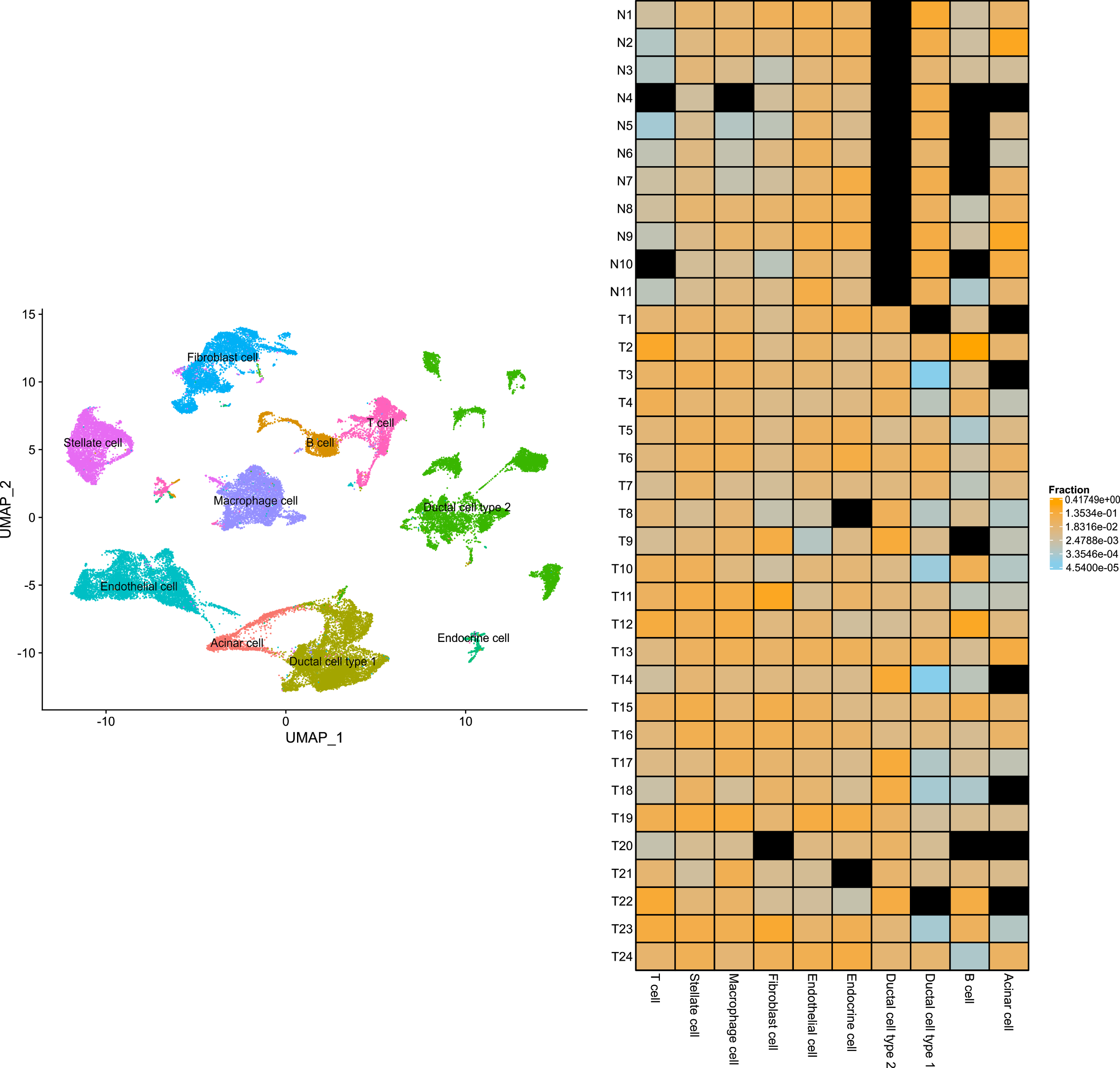

**Fig. S1.** Left: UMAP plot of all cell types in PDAC and non-PDAC samples in Peng et. al. Right: Fraction of cells from each cell type present in each non-PDAC (samples N1 to N11) and PDAC samples (T1 to T24). Squares filled with black represent zeroes.

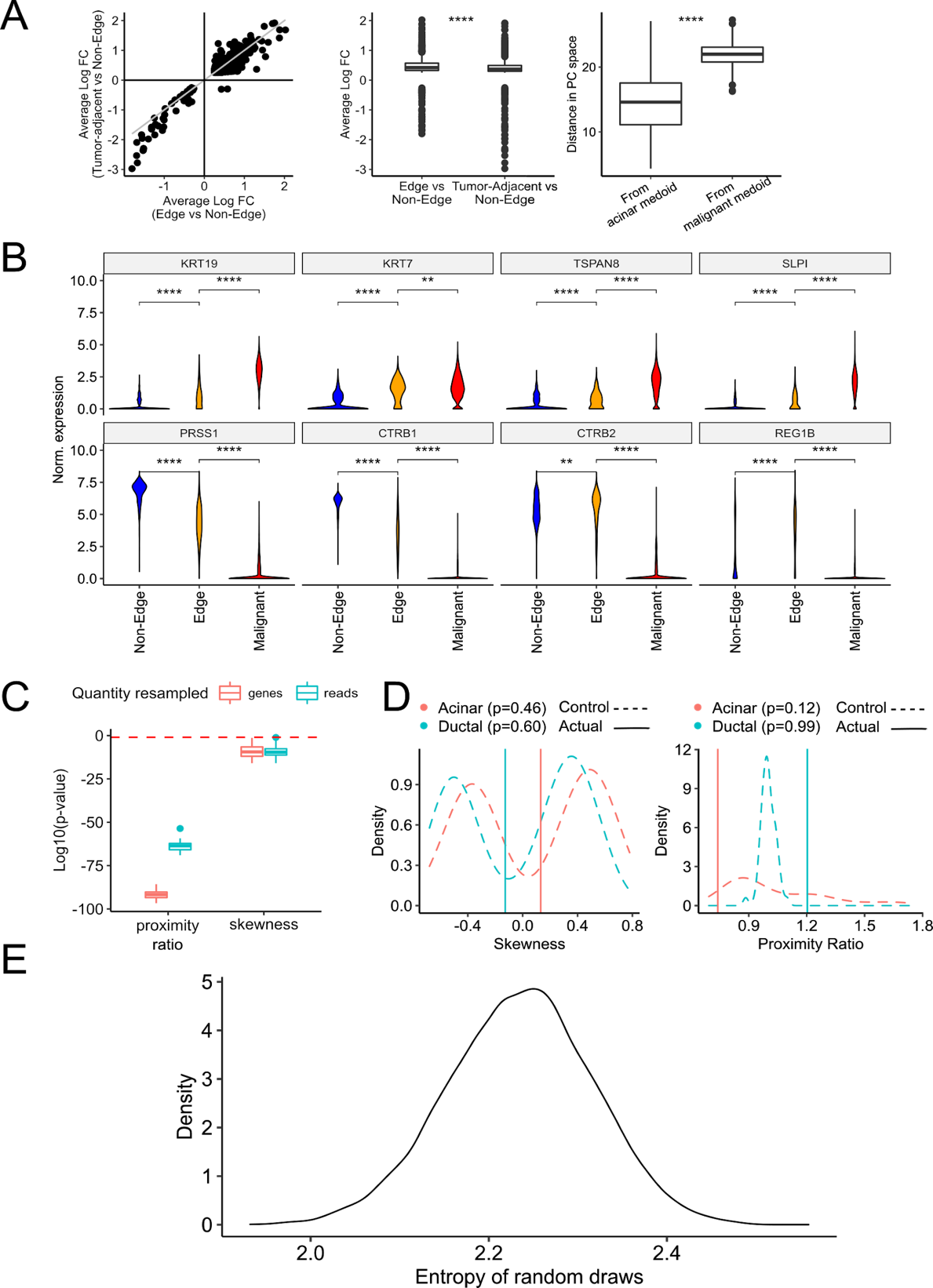

**Fig. S2**. (A) Left: Log-fold changes of differentially expressed genes between edge and non-edge acinar cells (x-axis) and between tumor-adjacent and non-tumor adjacent acinar cells (y-axis). Middle: The log-fold changes from the plot on the left visualized as a box-plot. Right: Distance of tumor-adjacent acinar cells from the acinar cluster medoid (left) and malignant cluster medoid (right). (B) Expression of marker genes of malignant ductal cells (top row) and acinar cells (bottom row). (C) P-values of skewness and proximity ratio tests after 100 re-samples of genes (left) and reads (right) from edge acinar cells. (D) Skewness and proximity ratio distributions of control and actual acinar and ductal cells from trajectory analysis-based pipeline described in Section S2. (E) Distribution of entropy values obtained from random draws of edge cells from multiple samples containing at least a single acinar cell, where each random draw generates a distribution of the fraction of edge cells arising from a sample.

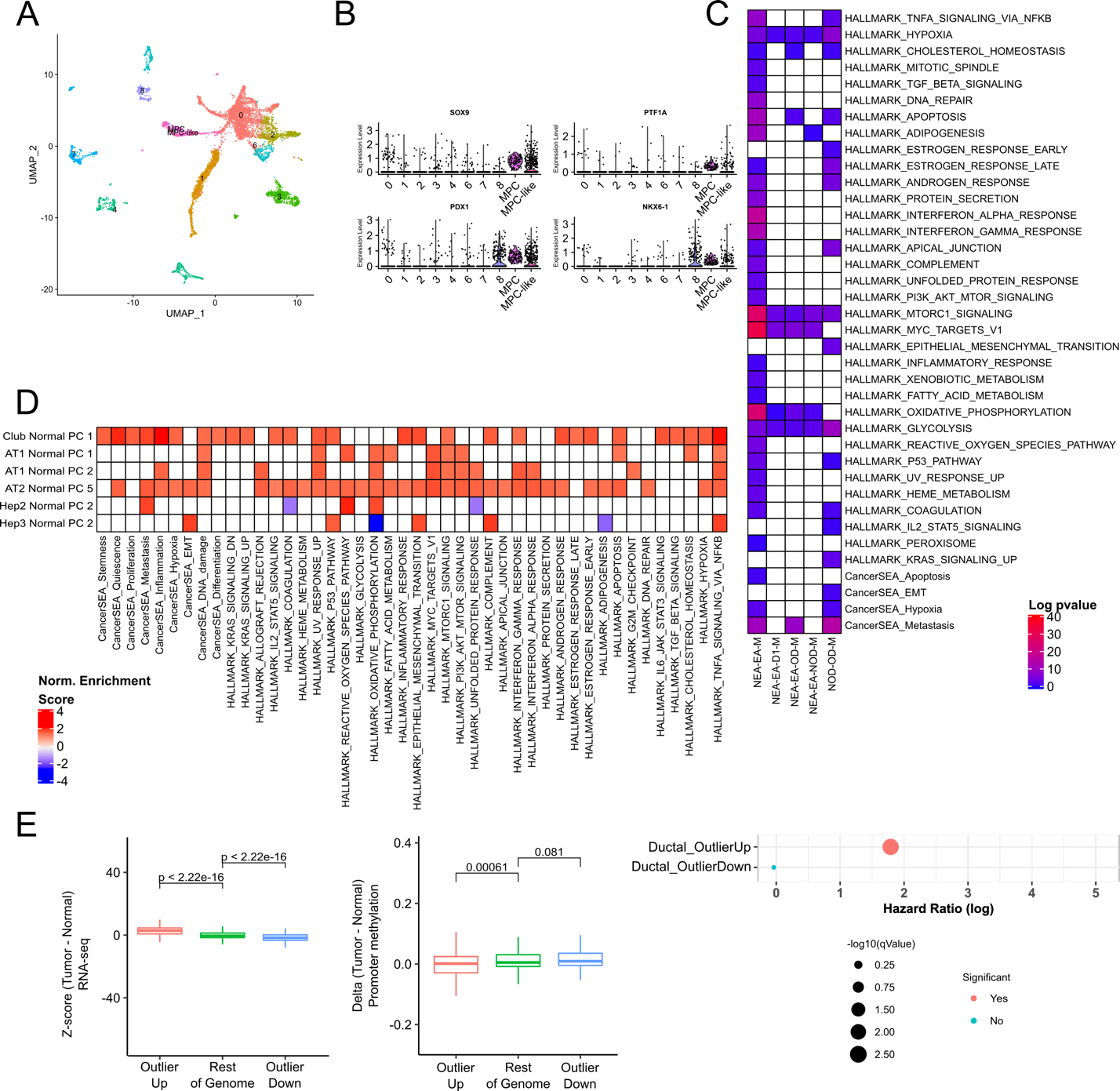

**Fig. S3.** (A) UMAP plot of clusters in fetal pancreas scRNA-seq data. (B) Expression of multipotent progenitor marker genes among clusters in fetal pancreas scRNA-seq data. X-axis represents the cluster Ids. (C) Pathways enriched among genes monotonically increasing in expression among five state transitions described in Fig. 2D i.e. Non-Edge -> Edge Acinar -> Malignant, Non-Edge Acinar -> Edge Acinar -> All Ductal -> Malignant, Non-Edge -> Edge Acinar -> Non-Outlier Ductal -> Malignant and Non-Outlier Ductal -> Outlier Ductal -> Malignant. (D) Hallmark and CancerSEA gene sets enriched amongst genes up-regulated in edge-like cells in lung and liver tissues. (E) Gene expression Z-scores (left column) and promoter methylation Z-scores (middle column) of Outlier-Up and Outlier-Down genes from TCGA data. The right column shows hazard ratios of the filtered Outlier-Up and Outlier-Down gene sets.
